# TOBF1 modulates mouse embryonic stem cell fate through co-transcriptional regulation of alternative splicing

**DOI:** 10.1101/2023.01.03.522557

**Authors:** Meghali Aich, Asgar Hussain Ansari, Li Ding, Vytautas Iesmantavicius, Deepanjan Paul, Chunaram Choudhary, Souvik Maiti, Frank Buchholz, Debojyoti Chakraborty

## Abstract

Embryonic stem (ES) cells retain the ability to undergo lineage-specific differentiation that can eventually give rise to different cell types that constitute an organism. Although stem cell specific biological networks of transcription factors and epigenetic modifiers are well established, how the ES cell specific transcriptional and alternative splicing (AS) machinery regulate their expression has not been sufficiently explored. In this study, we show that the lncRNA associated protein TOBF1 regulates the co-transcriptional alternative splicing of transcripts necessary for maintaining stem cell identity in mouse ES cells. Overlaying information derived from TOBF1 chromatin occupancy, the distribution of its pluripotency-associated OCT-SOX binding motifs, and transcripts undergoing differential expression and alternative splicing upon its disruption unmasked local nuclear territories where these distinct events converge, ultimately leading to the maintenance of mouse ES cell identity.

## Introduction

Mouse ES cells (mESCs) are derived from the inner cell mass of a developing blastocyst and are ex-vivo equivalent to the epiblast lineage, thus sharing the same developmental potential. These cells are characterized by their unique ability for high capacity in vitro self-renewal and the conservation of developmental pluripotency to differentiate into any of the three embryonic lineages, Ectoderm, Endoderm and Mesoderm (1–3). As such mESCs possess clinical potential for cell-based therapy and regenerative medicine research (4). To harness the quality and applicability of mESCs, it is important to understand the fundamental properties and molecular mechanisms that govern this unique identity.

Various studies have established the role of POU5F1, SOX2, NANOG, KLF4 and ESRRB as the core transcription factors that regulate the pluripotency network of mESCs (5–7). Pluripotent cells have distinct chromatin structure enriched for active histone marks and hyperdynamic binding of structural proteins (8). They also have global hyperactive transcriptional states and enhanced epigenetic modifiers as compared to differentiated cells (9, 10). The interplay between the pluripotency network and external cues orchestrates the in vivo self-renewal and undifferentiated stem cell state. These concepts have emerged from genetic, biochemical and molecular studies of transcription factors, chromatin regulators, non-coding RNAs, mRNA splicing and translational control (11–13).

In pluripotent cells, RNA Binding Proteins (RBPs) are part of the extended regulatory network that dynamically dictate the establishment of a stem cell-like state or induction of differentiation. Among the first RBPs, Lin28 proteins were recognized as regulators of pluripotency and stem cell maintenance (14). At various levels of post-transcriptional control, RBPs influence the stem cell state including RNA modification (METTL3/14, ADAR), alternative polyadenylation (FIP1), alternative Splicing (RBFOX2, SON), nuclear export (THOC2/5), RNA stability (TRIM71, ZFP36L1, PUM1) and translation (ESRP1, L1TD1) (15–17).

Alternative Splicing (AS) is a distinct event in metazoan genomes by which different combinations of exonic splice sites in pre-mRNA are selected to generate varied structural and functional mRNA and protein molecules (18–20). DNA binding and transcription factors also multitask at the level of RNA to regulate AS. For example, NACC1 regulates the expression of other splicing factors such as MBNL1 and RBFOX1 which indirectly controls the ES cell specific AS events to maintain the stem cell behavior (21). Depletion of the Spliceosome associated factor, SON leads to loss of pluripotency and cell death due to mis-splicing of key pluripotency factors OCT4, PRDM14, MED24 (22). Other splicing factors including RBM9, ESRP1, SFRS2, MBNL1/2, HNRNPLL play specific roles in regulating pluripotency by stimulating the precise splicing of various transcripts thereby altering global transcriptome expression (23–26). Another splicing factor hnRNPK, mediating proliferation and maintenance of myoblasts profoundly affects the cell cycle patterns thus indicating an initiation of differentiation (27).

Studies have shown the role of microRNAs and long non-coding RNAs in transcriptional regulation and post-transcriptional monitoring roles to support the self-renewal and pluripotency in ESCs (28, 29). Advances in high-throughput CLIP-Seq and genome association studies have revealed that lncRNAs bind with one or more RNA Binding Proteins (RBPs) and coordinately regulate gene expression (30, 31). In one study, it was seen that close to 75% of 226 lncRNAs expressed in mouse ESCs have binding sites for at least one of 9 pluripotency-associated transcription factors (OCT4, SOX2, NANOG, c-MYC, n-MYC, KLF4, ZFX, SMAD, and TCF3) and upon knockdown of 11 pluripotency-associated transcription factors, 60% mESC-expressed lncRNAs showed significant downregulation (32).

Previously, through a genome wide esiRNA screen study we had identified 594 lncRNAs targets for loss-of-function in mouse cells, of which lncRNA *Panct1* was molecularly characterized to have a positive role in maintaining the ES cell identity (33). A subsequent report revealed that the lncRNA *Panct1* interacts with an X-Chromosome associated protein TOBF1 (A830080D01Rik/BCLAF3) and transiently localizes to genomic loci containing the OCT-SOX motif in a specific cell-cycle regulated manner. This RNA-protein association is indispensable for maintaining the mouse ES cells state (34). Interestingly, in a recent study using a genome scale CRISPR screen, knockout of TOBF1 (also called BCLAF3) was revealed to support the Wnt-independent self-renewal of gastric epithelial cells in mouse gastric epithelial organoid models (35). This suggests that TOBF1 might have distinct roles dependent on individual cell type and/or tissue of expression.

TOBF1 in mice is derived from X-Chromosome and belongs to the BCLAF1/THRAP3 family of proteins and is also known as BCLAF1 and THRAP3 family member 3 (BCLAF3, Uniprot ID A2AG58). THRAP3 in humans is a spliceosome component and a subunit of the TRAP complex that has been reported to play a role in mRNA decay and pre-mRNA splicing (36). BCLAF1 and THRAP3 are RNA processing factors that have a role in DNA damage response pathway and genomic stability and prevention of oncogenic transformation (37, 38). Except for its role in self renewal of gastric epithelial cells in mouse (35), TOBF1’s role has remained largely uncharacterized, particularly in embryonic stem cells.

## Results

### TOBF1 interactome shows enrichment of splicing regulators

Our earlier studies on TOBF1 had clearly demonstrated distinct hallmarks of its cellular localization: bright puncta in early G1 phase of the cell cycle coupled with an association with chromatin. Notably, these were sites where the lncRNA *Panct1* localized as well and marked transient yet defined territories of unknown function (Chakraborty *et al*, 2017). To precisely pinpoint the mode of action of TOBF1 in this phase (Methods), we first performed an affinity purification followed by mass spectrometry-based identification of the associated proteins upon TOBF1 pulldown from a BAC tagged TOBF1-GFP mES cell line (Methods). A total of 672 proteins were pulled down and the bait (TOBF1-GFP) showed the highest enrichment (>20 peptides per replicate) as expected. In order to identify the potential interactors of TOBF1, we set up a stringent cutoff (-log T-test p value > 2) and further shortlisted potential interactors based on their presence in at least 2 out of 3 control pulldowns. Finally, we identified 17 proteins with high confidence that were significantly enriched in the TOBF1-GFP pulldown samples and not detected in controls (Figure 1A, Supplementary file 1).

**Figure 1:**
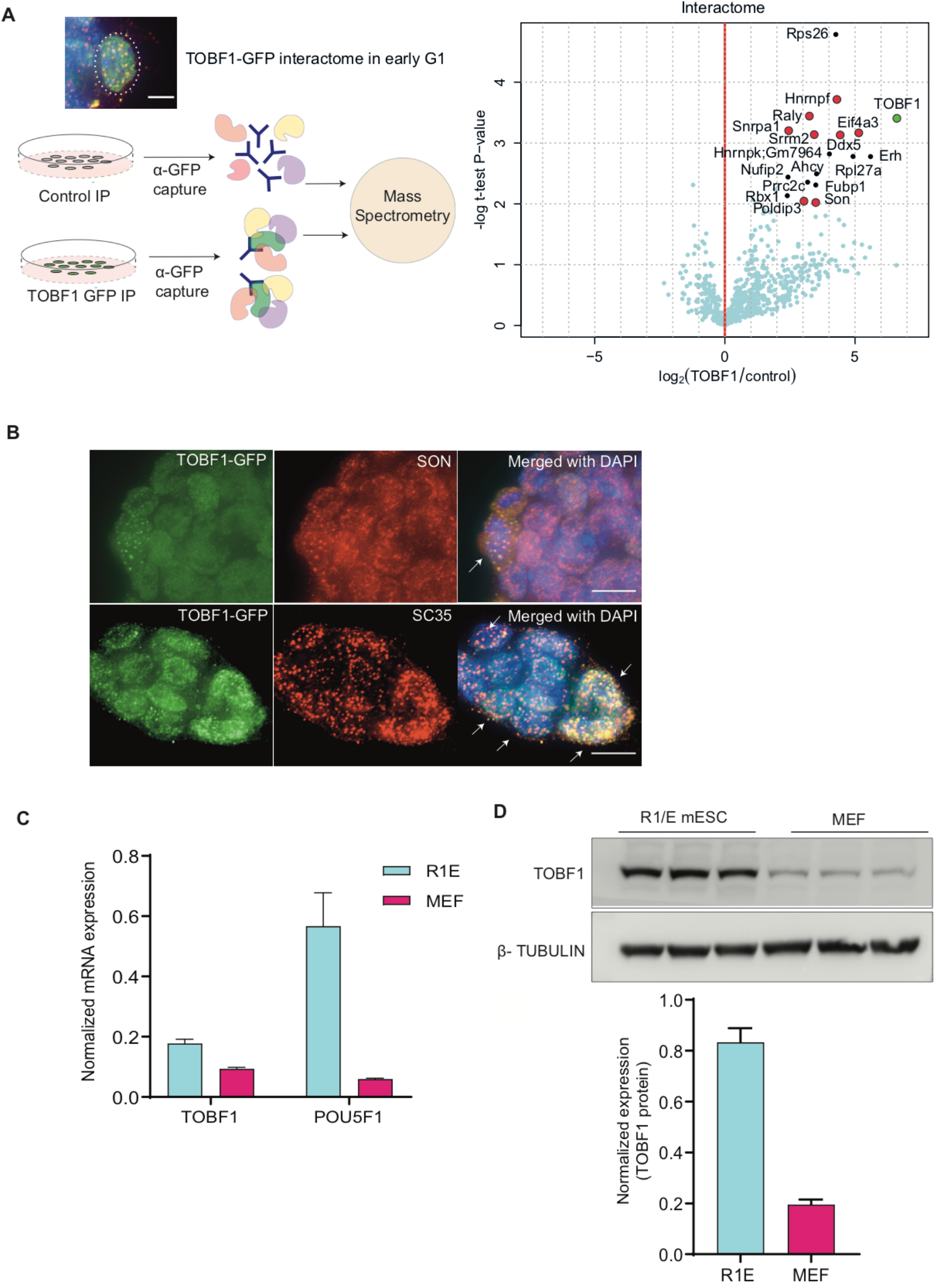
TOBF1 interactome in early G1 phase shows enrichment of splicing regulators. (A) Schematic showing TOBF1-GFP BAC-tagged cells form bright puncta in specific cell cycle phase. TOBF1-GFP in tagged cells is immunoprecipitated with anti-GFP antibody (control, R1/E mES cells) in the G1 phase through Demecolcine treatment and release (Methods). Mass spectrometry interactome profile showing interaction between TOBF1 and different mRNA processing and splicing proteins, a few of which are specific to stem cells (marked in red) with fold change (Log2) >=2 and p-value <0.05. (B) Representative Co-Immunofluorescence (Co-IF) images of combined TOBF1 (green) and SON (red) (upper panel) and TOBF1 (green) and SC35 (red) (lower panel), showing colocalization of TOBF1 and SON in TOBF1-GFP cells. Arrows mark colocalized spots. DNA is counterstained with DAPI and represented in blue. Scale bar 10μm. (C) qRT-PCR showing the steady state mRNA levels of Tobf1 and Pouf5f1 in R1E WT mES cells and MEFs, normalized with β-Actin. Error bars indicate the SD of three independent replicates. (D) Western blot showing expression of TOBF1 in R1E WT mES cells and MEFs as ~90KDa protein. β-Tubulin taken as loading control. Barplot with error bars showing SD of the quantification of the Western blot with three independent replicates.

Pathway analysis showed that 8 of the 17 proteins (47%) had roles in the processing and splicing of transcript isoforms by implementing various AS events like intron retention and exon skipping (Supplementary file 1). Some of these are well characterized regulators of specific mRNA processing, including SON, EIF4A3, SRRM2, ERH, RALY, hnRNP-F etc (39–42) (Figure 1A, Extended Fig1A). In addition, the interactome profile revealed ribosomal and ribonucleoproteins such as RPL27A, RPS26 and hnRNP-K that have been reported to affect the complex ribosomal biogenesis and mRNA stability pattern in ES-cell condition (43–45)) (Extended Figure 1A). Taken together, these results suggest that TOBF1 might be involved in the regulation of mRNA processing in mES cells.

Among the top TOBF1 interacting proteins, the presence of SON was particularly interesting since it has been implicated in the regulation of proper splicing of mRNA transcripts encoding pluripotency regulators in human ES cells (22). In a subsequent study, SON was shown to associate with BCLAF1 and THRAP3 (46), both of which belong to the same family as mouse TOBF1. Co-immunofluorescence (Co-IF) revealed that TOBF1 and SON colocalized (Pearson’s colocalization coefficient of ~0.6) in discrete puncta in ES cell nucleus (Figure 1B). Importantly, TOBF1 also showed colocalization with another canonical nuclear speckle marker and splicing regulator SC-35 (47) (Figure 1B) suggesting that the bright puncta in early G1 phase correspond to nuclear speckles (Pearson’s colocalization coefficient of ~0.75) potentially signifying a focal point for TOBF1’s RNA processing role.

Next, we performed co-immunoprecipitation of SON and TOBF1 and found that these proteins physically interact in mES cells (Extended Figure 1B). It is well known that mRNA processing and splicing factors often localize to the nuclear speckles (NSs) (48) and NSs are the hub for various mRNA post-transcriptional modifications. To validate that the TOBF1 interaction was specific to splicing associated factors present in nuclear speckles, we probed another spliceosome associated protein RPS26 and found that TOBF1 physically associates with it (Extended Figure 1B). These results establish that TOBF1 is indeed a bonafide member of the spliceosome associated proteome in the early G1 phase of the cell cycle in mES cells and the distinct TOBF1 puncta observed in the early G1 phase correlates with nuclear speckles.

To ensure that the BAC tagged TOBF1-GFP cell line retains the cytoarchitecture of the endogenous TOBF1 protein, we also validated the above observations with a commercially available TOBF1 antibody. We found that the TOBF1-GFP signal completely colocalized with the TOBF1 antibody signal (Extended Figure 1C). Importantly, in an untagged cell line (R1/E mES cells) too, the TOBF1 antibody displayed a mixture of bright puncta and diffused signal patterns as seen in the BAC-tagged cells (Extended Figure 1D). Together, this data shows that the TOBF1-GFP BAC tagged signal is a bonafide representation of the TOBF1 localization pattern in mES cells.

### High TOBF1 expression is a hallmark of mES cell pluripotency

Pluripotent cells have a complex regulatory network that distinctly differs from differentiated cells. The set of proteins required for the maintenance of an ES cell state are specific in their expression and function and as cells undergo differentiation, they are replaced by others (49, 50). Since esiRNA mediated transient knockdown of TOBF1 had previously shown a reduction in *Pou5f1* and *Klf4* mRNA levels (Chakraborty *et al*. 2017), both of which are signatures of pluripotent mES cells, we investigated if elevated expression of TOBF1 is a marker of undifferentiated stem cells. To this end we checked the expression of TOBF1 in R1E pluripotent mES cells and compared it to a mouse embryonic fibroblast (MEF) line which are differentiated and do not show pluripotent characteristics. We observed that both at the level of mRNA and protein levels TOBF1 expression in mES cells was higher (~2 fold at mRNA and ~4 fold at protein levels) than MEF cells suggesting that as cells exit from pluripotency, TOBF1 levels decline (Figures 1C-D). Next, we investigated the heterogeneity in TOBF1 levels across a small subset of mouse tissues and cell lines (Liver, Kidney, Lung, Heart, Prefrontal cortex, Reticular Cortex (RC), Hippocampus (HC), Hypothalamus (HT), Medulla Oblongata (OB), Neuroblastoma (N2A) and Murine Melanoma (B16) (Extended Figure 1E). Even within this subset, we found that TOBF1 expression was highest in mES cells further corroborating its identity as a bonafide ES cell marker. A recent report has suggested that TOBF1 is expressed in non-stem cells in mice and regulates gastric epithelial differentiation. This suggests that TOBF1 expression might be heterogeneous and have additional tissue specific roles (35).

**Extended figure 1:**
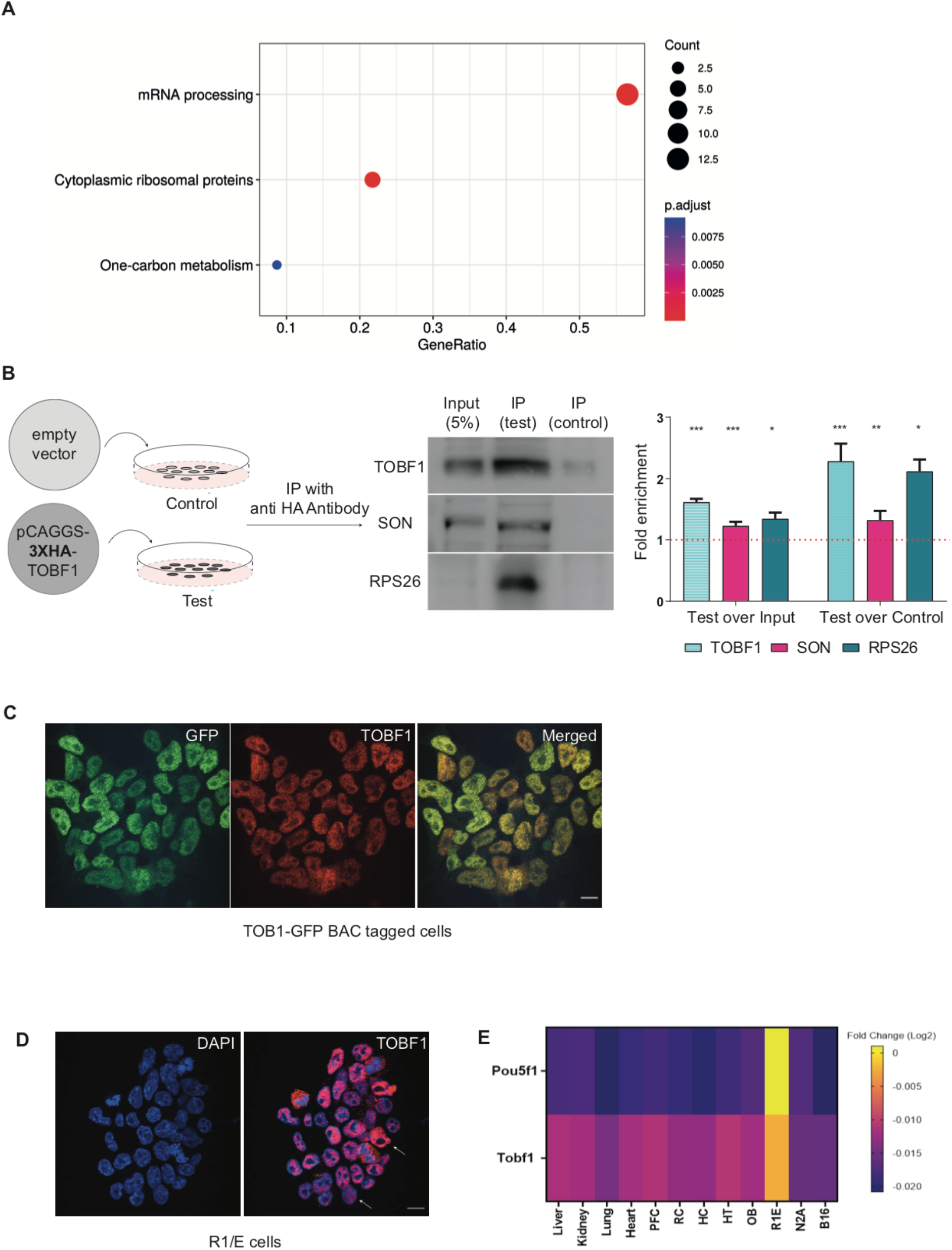
(A) Dot plot of wikipathways showing the pathway analysis of the proteins from the TOBF1 interactome depicts enrichment of pathways associated with mRNA processing and cytoplasmic ribosomal proteins, adjusted p-value 0.001-0.01. Dark red represents the high significant p-value and dot scaled with respect to the number of genes involved in pathways. (B) Schematic showing Immunoprecipitation (IP) in transient overexpression with 3XHA TOBF1 expressing plasmid system (Test) and R1E mES cells (control). IP was done using anti-HA antibody and representative western blot image showing enrichment of TOBF1 in IP-Test samples as compared to Input (5%) and IP-Control. SON and RPS26 enrichment are also observed in TOBF1 pull down samples as compared to control. Bar plot showing the quantification of protein enrichment. Error bars indicate the SD of three independent replicates. *p<0.05, **p<0.001, ***p<0.0001 (Student’s t-test). (C) Representative Co-Immunofluorescence image of TOBF1 (using anti-TOBF1 ab, shown in red) and GFP (shown in green) showing co-localization of TOBF1 and GFP (shown as yellow dots) in TOBF1-GFP BAC tagged mES cell line. Scale bar 10 μm. (D) Representative image of TOBF1 signals (shown in red) in R1/E mES cells using TOBF1 antibody that displayed the mixture of bright puncta and diffused signal patterns as seen in the BAC-tagged cells. DNA is counterstained with DAPI. Scale bar 10 μm. (E) Heatmap showing the absolute steady state mRNA levels of Tobf1 in different mouse tissues in small subset of mouse tissues and cell lines (Liver, Kidney, Lung, Heart, Prefrontal cortex, Reticular Cortex (RC), Hippocampus (HC), Hypothalamus (HT), Medulla Oblongata (OB), Neuroblastoma (N2A) and Murine Melanoma (B16). POUF51 is considered as a positive control for mES cells. All values are normalized to β-Actin.

### TOBF1 deficient mES cells exit from pluripotent state

In our previous study, we had identified that TOBF1 interacts with a lncRNA Panct1 and forms bright nuclear puncta at G1 cell cycle phase. Although this localization pattern could be extrapolated to chromosomal territories both by electron microscopy and Chromatin Immunoprecipitation (ChIP), our results did not establish the functional role of TOBF1 as a distinct regulator of transcription.

Since TOBF1 has a prominent cell cycle specific localization pattern we next inquired if this was influenced by gross changes in TOBF1 levels across the cell cycle. We sorted R1/E mES cells using FUCCI (Fluorescent, Ubiquitination-based cell cycle indicator) system for distinguishing the G1 and non G1 phases of the cell cycle (51). We noticed a small (~1.5 fold) increase in TOBF1 mRNA levels in early G1 phase as compared to non G1 phase cells (Figure 2A) suggesting that TOBF1 levels might undergo a low level of shuttling between its G1 versus non G1 expression patterns.

**Figure 2:**
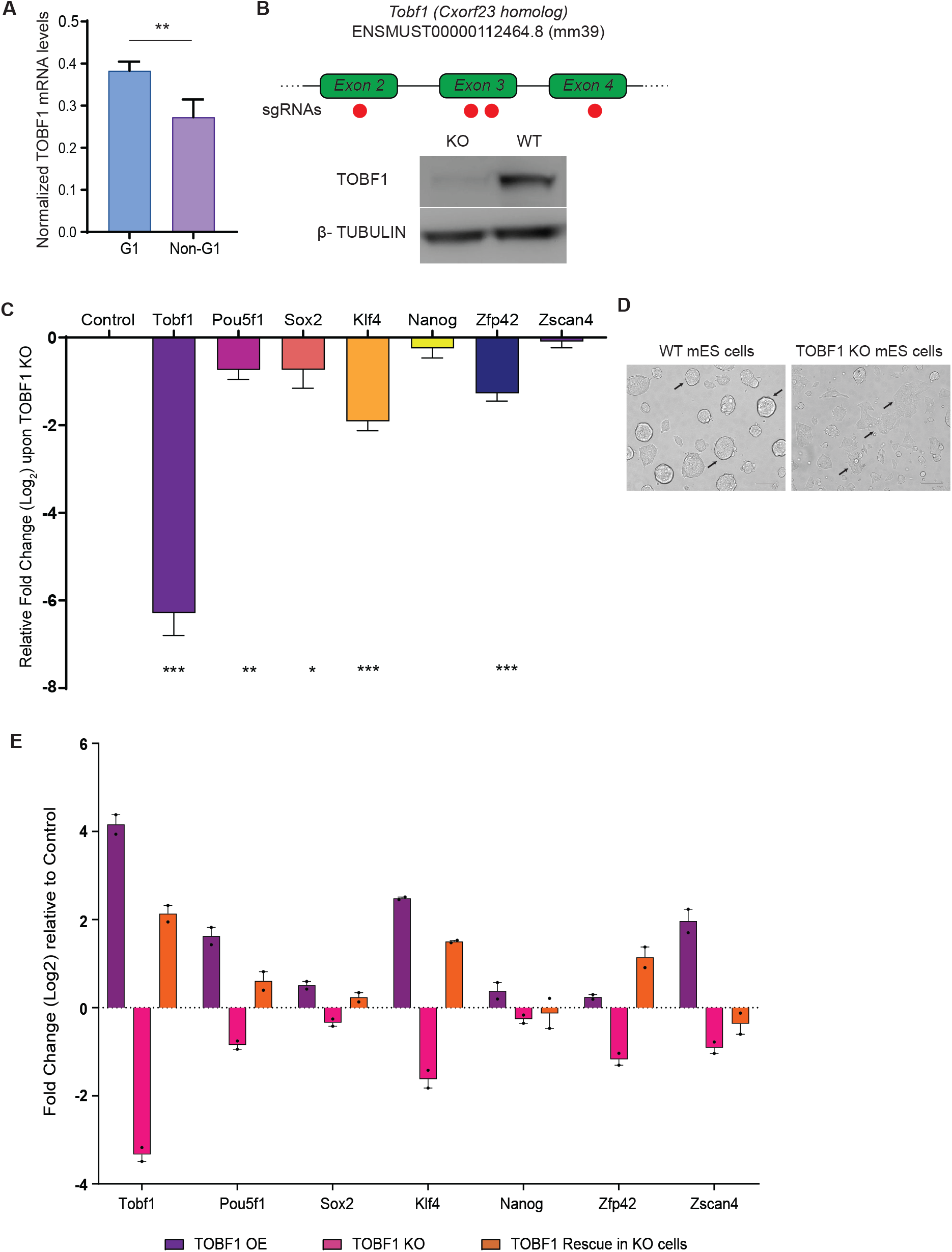
TOBF1 Knockout cells mES cells exit from the pluripotent state. (A) Bar plot showing absolute steady state mRNA levels of Tobf1 in G1 and non-G1 sorted cells using ES-FUCCI system. Tobf1 values are normalized with β-Actin. Error bars indicate the SD of three independent replicates. *p<0.05, **p<0.001, ***p<0.0001 (Student’s t-test). (B) Schematic diagram of the dual sgRNA based CRISPR targeting of exons 2, 3 and 4 of Tobf1 gene. Representative image of western blot showing TOBF1 protein expression in R1E WT mES cells and TOBF1 KO cells as ~90KDa protein. β-Tubulin taken as loading control. Bar plot with error bars showing SD of the quantification of the Western blot with three independent replicates. ***p<0.0001 (Student’s t-test). (C) qRT-PCR showing relative fold change (Log2) of mRNA levels of different pluripotency markers (*Pou5f1, Sox2, Klf4, Nanog, Zfp42, Zscan4*) upon TOBF1 knockout as compared to R1/E WT control cells. All values normalized with β-Actin. Error bars indicate the SD of three independent replicates. *p<0.05, **p<0.001, ***p<0.0001 (Student’s t-test). (D) Representative grayscale light microscope images showing distinct morphological differences between R1/E WT cells and TOBF1 KO cells. Undifferentiated R1/E WT mES cells form round intact colonies but TOBF1 KO cells show more spread-out flat morphology. Scale bar 100μm. (E) qRT-PCR showing relative mRNA fold levels of Tobf1 and other pluripotency markers (*Pou5f1, Sox2, Klf4, Nanog, Zfp42, Zscan4*) in TOBF1 overexpression, TOBF1 knockout and TOBF1 rescue conditions as compared to R1/E WT cells. All values normalized with β-Actin. Error bars indicate the SD of two independent replicates.

We next turned our attention to the possible role of TOBF1 in regulating ES cell state through perturbation experiments. We adopted two CRISPR based approaches to either knockdown (CRISPR interference, CRISPRi) (52); (53) or knockout (CRISPR based dual sgRNA mediated deletion) (54) TOBF1 in R1/E WT mES cells. For TOBF1 knockdown, we used a dFnCas9 fused with Krüppel-associated box (KRAB) domain (55–57) along with sgRNAs targeting the promoter of TOBF1 locus (Extended Figure 2A). We observed that 48hrs after transfection of CRISPRi constructs, a ~70% reduction in TOBF1 mRNA levels were seen. Importantly, a concomitant reduction in classical pluripotency and pluripotency associated markers such as *Pou5f1, Sox2, Klf4, Nanog, Zfp42* and *Zscan4* was observed suggesting that these cells were exiting from their pluripotent state (Extended Figure 2B). This observation was similar to endoribonuclease prepared siRNA (esiRNA) mediated knockdown studies where TOBF1 mRNA was targeted and similar reduction in *Pou5f1* and *Klf4* mRNA levels were seen. Both these results confirm that TOBF1 perturbation impairs the expression of regulators of pluripotency in mESCs.

Having established the connection between TOBF1 mRNA levels and markers of mES cell pluripotency, we next proceeded to investigate mES cells lacking TOBF. To this end, we generated CRISPR mediated stable TOBF1 knockout in R1/E mESCs using single guide RNAs (sgRNAs) targeting exons 2, 3 and 4. We observed efficient knockout of TOBF1 expression both at mRNA and protein levels (Figure 2B). Similar to the knockdown experiments, upon TOBF1 knockout, the expression of the key pluripotency regulators such as *Pou5f1, Sox2, Klf4, Nanog, Zfp42* and *Zscan4* were all downregulated (Figure 2C). As a consequence of exit from pluripotency, TOBF1 KO mES cells displayed a flattened morphological phenotype comparable to differentiated cells (Figure 2D). Taken together with our earlier observations, we conclusively show that TOBF1 perturbation through transcriptional repression of the gene promoter, genetic knockout of the gene, or RNA interference all result in an exit from pluripotency.

Since TOBF1 KO cells showed reduced expression of pluripotency associated genes, next we investigated if this phenotype could be rescued by exogenous overexpression of TOBF1. Upon overexpressing TOBF1 through a constitutive promoter, we observed that apart from *Nanog* and *Zscan4*, the expression of all other pluripotency genes (*Pou5f1, Sox2*, *Klf4* and *Zfp42*) switched from a downregulated state to an upregulated state (Figure 2E). This further strengthened the association of TOBF1 with the core pluripotency network in mES cells and TOBF1 overexpression shows a similar phenotype as rescue but with higher fold change. We then sought to understand if modulating TOBF1 brings any changes in the cell cycle pattern of the ES cells where exit from pluripotency results in reduction in the S phase population (58, 59), (60); (61). Interestingly, upon TOBF1 rescue, the proportion of S phase cells increased(Extended Figure 2C) suggesting that the exogenous expression of TOBF1 in KO cells could rescue its phenotype and in turn induce self-renewal capacity.

**Extended Figure 2:**
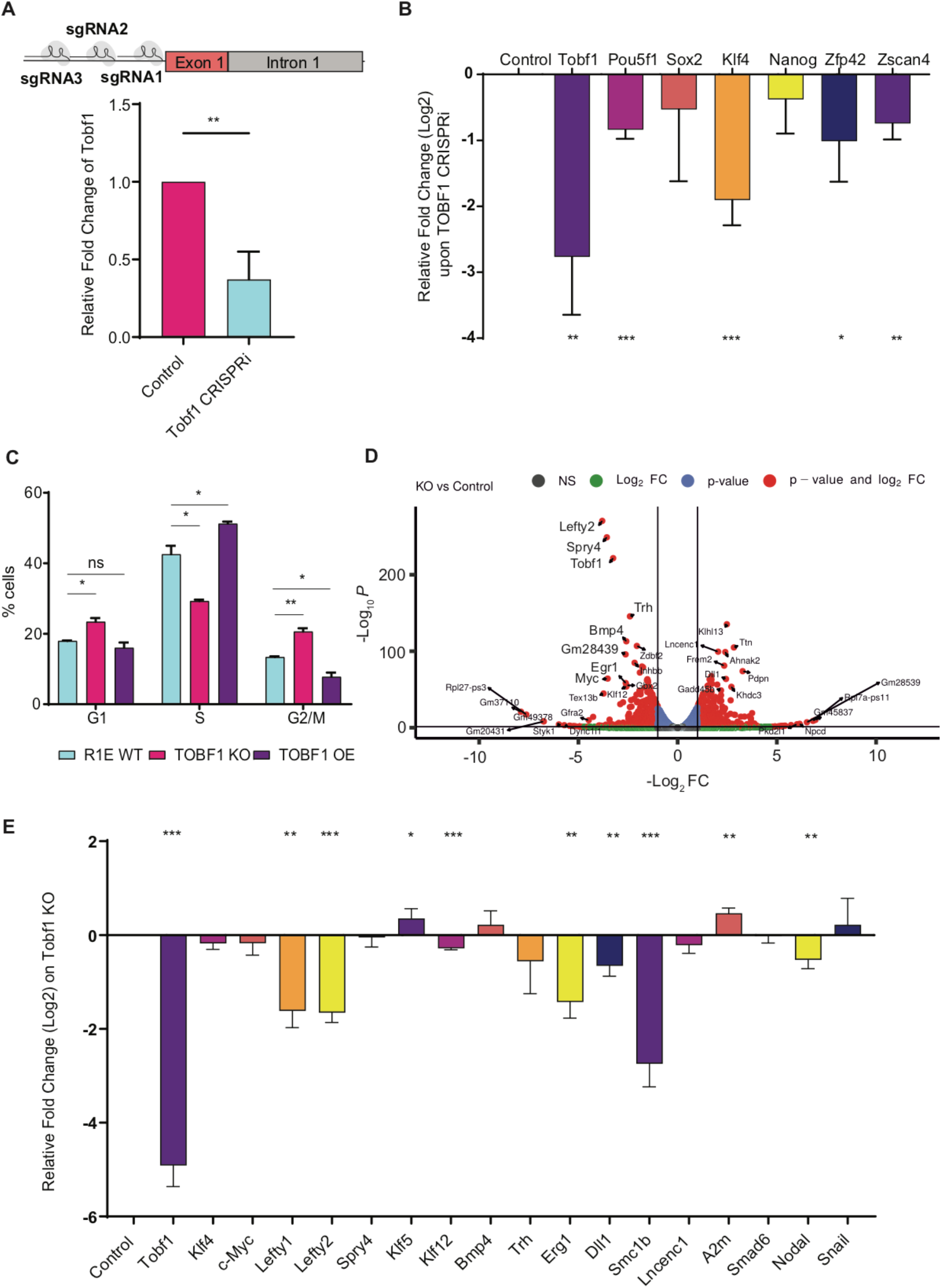
(A) Schematic diagram showing the sgRNAs targeting the promoter of Tobf1 gene for CRISPRi based gene downregulation. qRT-PCR showing Tobf1 mRNA levels in Control and TOBF1 knockdown samples. Values normalized with β-Actin. Error bars indicate the SD of three independent replicates. *p<0.05, **p<0.001, ***p<0.0001 (Student’s t-test). (B) Bar plot showing the relative steady state mRNA levels of different pluripotency markers in TOBF1 knockdown samples as compared to R1/E WT (control) cells. All values normalized with β-Actin. Error bars indicate the SD of three independent replicates. *p<0.05, **p<0.001, ***p<0.0001 (Student’s t-test). (C) Bar plot showing distribution of cells in G1, S, G2/M phase of cell cycle in R1/E WT, TOBF1 knockout (KO) and TOBF1 overexpressed (OE) cells. Error bars indicate the SD of three independent replicates. *p<0.05, **p<0.001, ***p<0.0001. (D) Volcano plot showing differential gene expression from RNA-Seq analysis of TOBF1 KO and R1/E WT cells. The horizontal line corresponds to a significance value of Log10 p-value <0.05. The two vertical lines bound the minimal fold change (Log2 >=|1|) for the most differentially expressed genes. (E) qRT-PCR showing the relative mRNA fold change (Log2) for the validation of the top 15 differentially expressed genes from RNA-Seq analysis in TOBF1 KO samples as compared to WT. All values normalized with β-Actin. Error bars indicate the SD of three independent replicates. *p<0.05, **p<0.001, ***p<0.0001 (Student’s t-test).

### TOBF1 perturbation affects global transcript levels associated with pluripotency and lineage commitment

Having established the connection between TOBF1 and mES cell pluripotency, we investigated how TOBF1 KO influenced the genome-wide transcriptomic profile through total RNA-Sequencing (RNA-Seq). Importantly, since the nuclear puncta phenotype and the association with splicing regulators were prominently observed in G1 cell cycle sorted TOBF1 KO cells, we focused on this phase for studying transcriptomic changes.

We used the FUCCI system to sort G1 specific mES population in TOBF1 KO cells and observed that 4644 genes (2253 Up-regulated and 2391 Down-regulated) were differentially expressed between TOBF1 KO and WT with FDR < 0.05 (Supplementary file 2). We performed pathway analysis of these genes and found that the top two pathways enriched (p<0.01) were ‘mRNA processing’ and ‘mechanisms associated with pluripotency’ (Figure 3A, Supplementary file 3). Although it was expected that pluripotency genes would be differentially expressed upon TOBF1 KO based on our RT-qPCR results, mRNA processing genes being affected signified a hitherto unexplained role of the protein which we subsequently investigated. Interestingly, among the topmost genes that showed the most significant fold reduction were *Klf4, Lefty2, Lefty1, c-Myc, Spry4, Sox2, Klf5, Klf12, Bmp4* all having known roles in regulating mES cell identity (62–73). In contrast, lineage commitment factors such as *Dll1* and TGF-β signaling factors *Smad6* (74)8; (75) were significantly upregulated (Extended Figure 2D). We selected a subset of the top differentially expressed transcripts from the TOBF1 RNA-seq results (*Klf4*, *c-Myc, Lefty1, Lefty2, Spry4, Klf5, Klf12, Bmp4, Trh, Egr1, Dll1, Smc1b, lncenc1, A2m, Smad6*) and successfully validated several of them using quantitative real time PCR (RT-qPCR). In addition, we found *Nodal*, another bonafide pluripotency (76) marker down-regulated upon TOBF1 KO (Extended Figure 2E). Through these results we concluded that TOBF1 KO leads to global transcriptome wide changes that cumulatively result in an exit from pluripotency.

**Figure 3:**
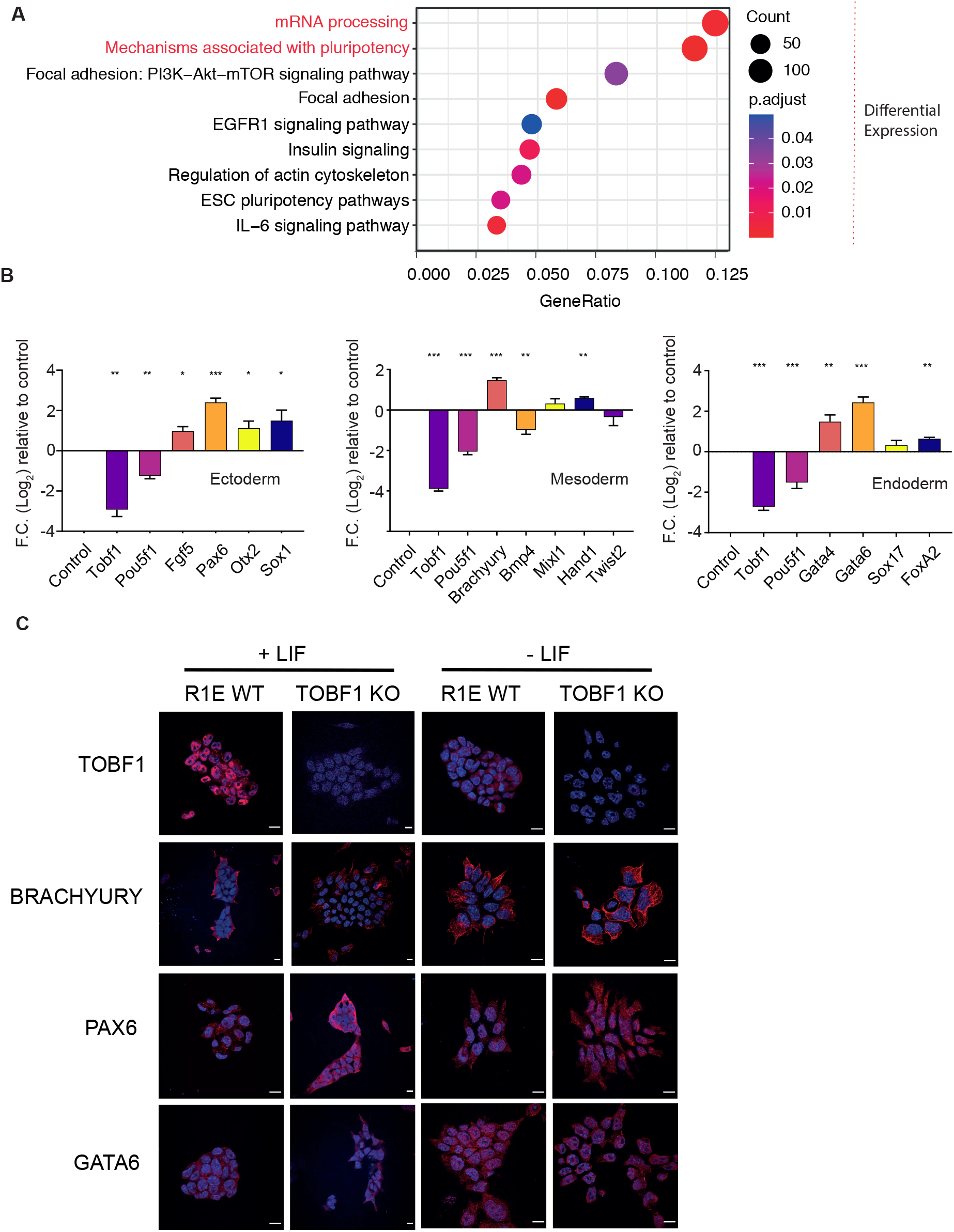
TOBF1 perturbation affects global transcript levels associated with pluripotency and lineage commitment. (A) Dot plot showing the enrichment of the top pathways of the differentially expressed genes from RNA-Seq. Dark red represent the high significant p-value and dot scaled with respect to number of genes involved in pathways. (B) qRT-PCR showing the relative mRNA fold levels of ectodermal (*Fgf5, Pax6, Sox1, Otx2*), mesodermal (*Brachyury, Bmp4, Mixl1, Hand1, Twist2*) and endodermal (*Gata4, Gata6, Sox17, FoxA2*) lineage markers in TOBF1 KO as compared to R1/E WT cells when these cells are cultured in defined trilineage differentiation media. Pou5f1 is taken as a negative marker for differentiation. All values normalized with β-Actin. Error bars indicate the SD of three independent replicates. *p<0.05, **p<0.001, ***p<0.0001 (Student’s t-test). (C) Representative images of Immunofluorescence (IF) showing expression of TOBF1, BRACHYURY, PAX6 and GATA6 (shown in red) in R1/E WT mES cells and TOBF1 KO cells cultured in regular media (+LIF, 20%FBS) and spontaneous differentiation media (-LIF, 10%FBS). DNA is counterstained with DAPI. Whiskers indicate positive and negative deviation from mean (n=40 cells). Scale bar 10 μm.

Next we investigated if differentiated TOBF1 KO cells have an increased expression of lineage commitment factors. To this end, we differentiated TOBF1 KO ESCs in a trilineage differentiation media and observed that markers corresponding to all three germ layers (*Fgf5, Pax6, Otx2* and *Sox1* for ectoderm; *Brachyury, Mixl1, Hand1* and *Twist2* for mesoderm and *Gata4, Gata6, Sox17* and *FoxA2* for endoderm) (77–86) showed enhanced expression (Figure 3B). These results suggested that TOBF1 KO primes ES cells to enter into a differentiated state.

As a corollary, we also tested whether subjecting TOBF1 KO cells to random differentiation by removal of LIF enhanced the expression of lineage commitment markers in these cells. Using qRT-PCR we confirmed that lineage markers corresponding to all three germ layers (*Brachyury, Hand1, Twist2* and *Mixl1* for ectoderm, *Otx2, Sox1* and *Pax6* for ectoderm and *Gata4, Gata6, Sox17* and *FoxA2* for endoderm) were overexpressed over control (Extended Figure 3A). This was further confirmed at the protein expression level using Immunofluorescence for PAX6 (ectoderm), Brachyury (mesoderm) and GATA6 (endoderm) (Figure 3C and Extended Figure 3B). Interestingly, we observed that upon spontaneous differentiation, TOBF1 KO leads to a comparatively higher expression of mesodermal markers suggesting that genes involved in the mesodermal lineage might be strongly affected upon TOBF1 KO. In concordance with this, among the RNAseq hits, both *Lefty1* and *Lefty2* both of which are prominent members of the TGF-β pathway and regulate mesoderm formation in vertebrates (87, 88) emerged as highly differentially expressed upon TOBF1 KO (Extended Figure 2D-E).

**Extended Figure 3:**
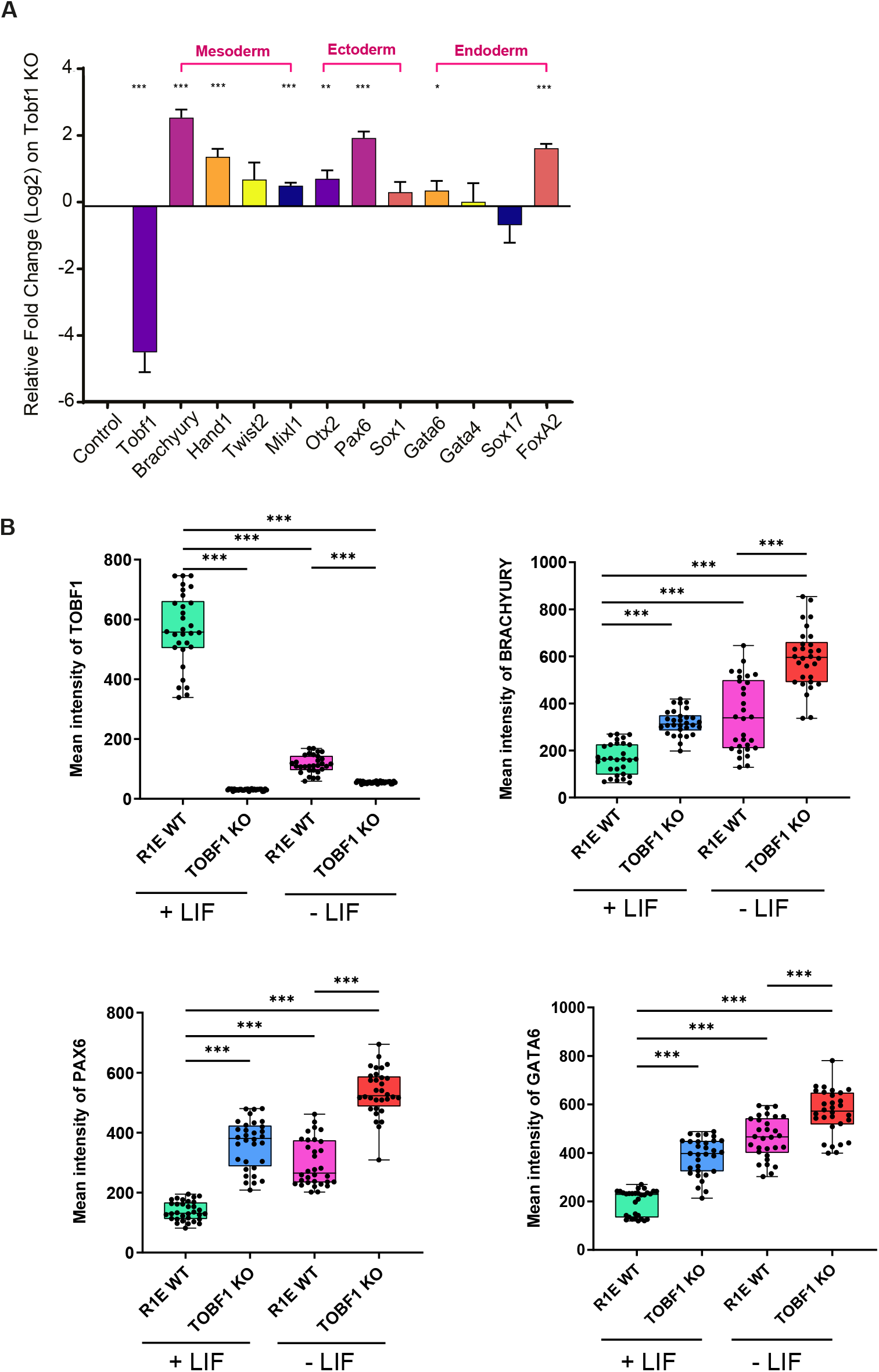
TOBF1 KO cells show enhanced expression of lineage committed markers. (A) qRT-PCR showing relative mRNA fold levels of different lineage markers as ectodermal (*Fgf5, Pax6, Sox1, Otx2*), mesodermal (*Brachyury, Bmp4, Mixl1, Hand1, Twist2*) and endodermal (*Gata4, Gata6, Sox17, FoxA2*) in TOBF1 KO and R1/E WT control cells when these cells are cultured in spontaneous differentiation media for 7 days. All values normalized with β-Actin. Error bars indicate the SD of three independent replicates. *p<0.05, **p<0.001, ***p<0.0001 (Student’s t-test). (B) Box plot showing the quantification of the Immunofluorescence (IF) of TOBF1, BRACHYURY, PAX6, GATA6 expression (shown in red) in R1/E WT cells and TOBF1 KO cells when cultured in normal media (+LIF, 20% FBS) and spontaneous differentiating media (-LIF, 10%FBS) for 7 days.

### Absence of TOBF1 alters mRNA splicing patterns in mES cells

Pathway analysis of the differentially expressed transcripts upon TOBF1 KO revealed that genes responsible for mRNA processing were even more significantly altered than pluripotency genes. Although mRNA processing involves a multitude of pathways including capping, methylation, splicing, 3’ end processing etc. we decided to focus on the role of TOBF1 on alternative splicing (AS) since interactome studies had already established its link with splicing factors (Figure 1A).

TOBF1 knockout cells showed 11235 events (from 5502 genes) (DPSI > |0.10|) that had undergone differential alternative splicing (Extended Figure 4A, Supplementary file 4). Strikingly, the top two cellular pathways which these events contributed to (based on the number of enriched genes) were ‘mRNA processing’ and ‘mechanisms associated with pluripotency’, both of which were enriched among differentially expressed transcripts as well (Figure 4A, Supplementary file 5). Apart from these two pathways, EGFR1 signaling and Insulin signaling also featured in the top pathways enriched among differentially spliced transcripts.

**Figure 4:**
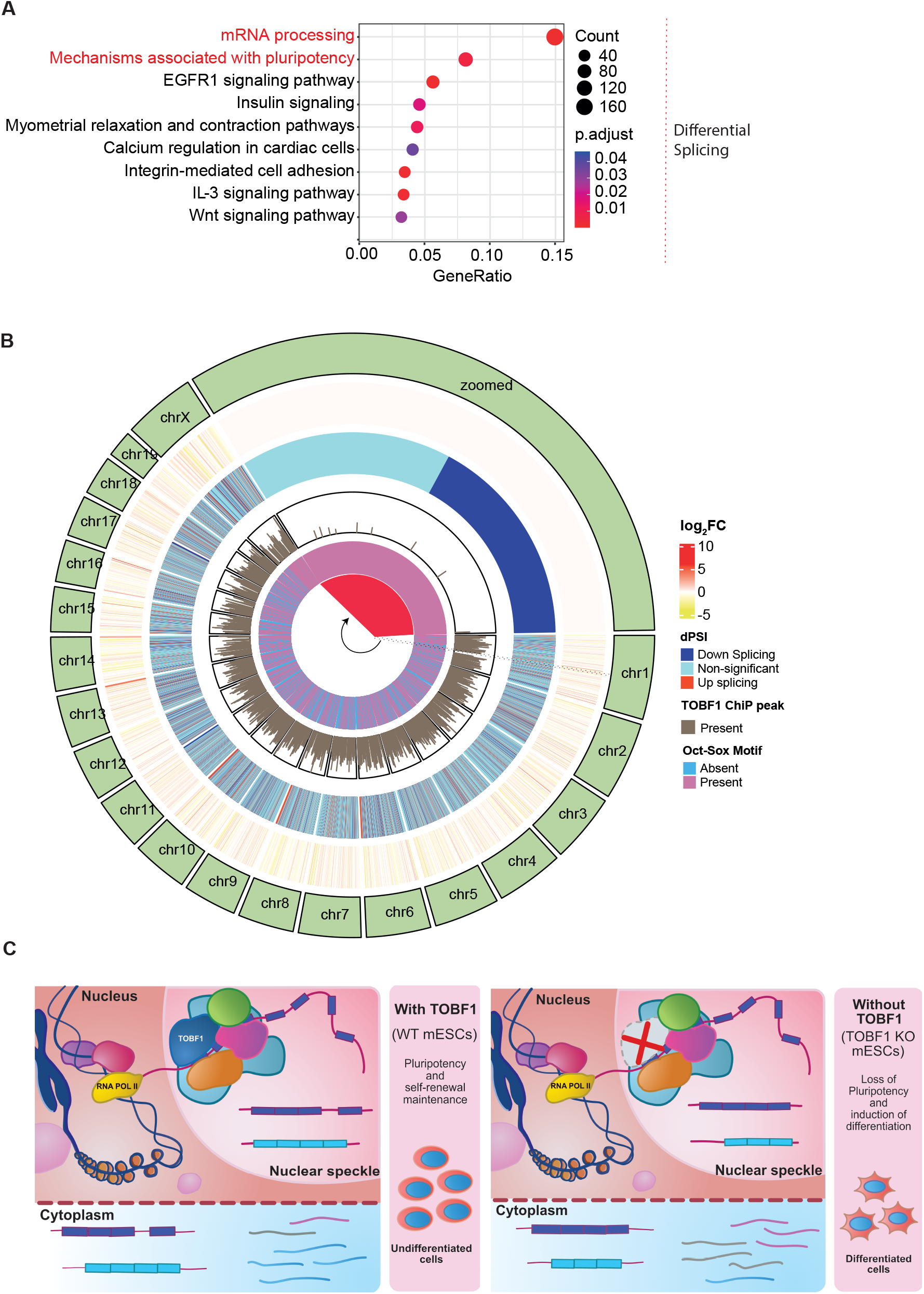
Absence of TOBF1 alters mRNA splicing patterns in mES cells. (A) Dot plot of wikipathways showing the enrichment of the top pathways of the differentially alternatively spliced events from RNA-Seq. Dark red represents the high significant p-value and dot scaled with respect to the number of genes involved in pathways. (B) Circos plot showing TOBF1 chromatin occupancy is strongly correlated with sites of CTS and also these events strongly correlated with the presence of the OCT-SOX motifs on DNA.

The enrichment of common pathways related to mRNA processing and ESC pluripotency suggested that there was a potential overlap of transcripts between the two pathways. Indeed, 27% of transcripts (68 out of 249) involved in mRNA processing (Extended Figure 4B) and 23% of transcripts (43 out of 185) involved in ESC pluripotency (Extended Figure 4C) were both significantly differentially expressed and differentially spliced. Strikingly, the transcripts that were alternatively spliced included the well-known regulators of alternative splicing themselves, namely NCL and SRSF1. To this end, we validated the alternative splicing of both the transcripts using qRT-PCR where *Srsf1* AS isoform was strongly downregulated (upto ~91 %) and *Ncl* AS isoform showed a robust upregulation (upto ~ 11.8 fold) (Extended Figure 4D).

The presence of common transcripts that were both alternatively spliced and differentially expressed appeared surprising considering that transcription regulation and alternative splicing of the same gene proceeds through different mechanisms (89–91). Further, the consequence of TOBF1 perturbation pluripotency appeared to be a cumulative effect of multiple altered ES cell related pathways that all contributed to the exit from undifferentiated state. For example, the highly significant AS events included KLF4, an important pluripotency master regulator, JARID2, a Polycomb Repressive Complex 2 (PRC2) component which regulates ES cell specific transcriptional signatures (92, 93) and CTCF a multifunctional protein that mediates long range interactions modulating pluripotency (94, 95). In addition to these, several other factors which are known to be important regulators of ES cell pluripotency such as ESRRB, HDAC1 and 4, DNMT3A etc which function through multiple distinct pathways (96–101) were also alternatively spliced (Extended file 1).

In a recent study in human ESCs, it was found that hnRNP(H/F) mediated AS of *Tcf3* lead to distinct isoforms that eventually led to hESC differentiation (102). This report could pinpoint a distinct AS event which appears to affect the downstream E-cadherin mediated exit from pluripotency. In contrast TOBF1 KO AS events are highly diverse in nature and span multiple pathways ranging from local transcriptional events to long range chromatin modifications. Importantly the differentially expressed genes upon TOBF1 KO also include a multitude of pluripotency pathways encompassing core transcription factors as well as well-known regulators of lineage commitment.

### TOBF1 forms discrete co-transcriptional splicing (CTS) hubs at sites of OCT-SOX motifs

Considering that TOBF1 is a global regulator of splicing and transcription, we considered its punctate localization in mESC nuclei during the early G1 phase as a possible hub for co-transcriptional splicing (CTS). This process which happens in close spatially and temporally associated territories have been reported in multiple systems although its role in ES cells have not been reported to our knowledge.

Our earlier studies on the association of *Panct1* and TOBF1 at OCT-SOX motifs had suggested the possible binding of TOBF1 to chromatin in the bright nuclear puncta in early G1 cells (34). In this study we identified TOBF1 to be a component of the spliceosome complex whose perturbation led to global splicing changes. Importantly, this also led to transcriptional changes which finally culminated in loss of pluripotency. The three observations, namely DNA occupancy, splicing modulation and transcriptional regulation are all hallmarks of CTS and indicate the formation of local factories or hubs of such coordinated events which are potentially scaffolded through lncRNAs like *Panct1*. Incidentally, the highly abundant lncRNA *Malat1* is known to regulate AS events through localization to nuclear speckles and interaction with splicing factors through conserved motifs in its secondary structure (103, 104).

To investigate if the two TOBF1 events (DNA binding and CTS) are indeed associated, we overlaid the three datasets of chromatin occupancy, splicing events and differentially expressed transcripts upon KO across the whole mouse genome (Circos plot, Figure 4B, Extended sheet 4). We observed that TOBF1 chromatin occupancy was strongly correlated with sites of CTS. Moreover, both these events also strongly correlated with the presence of the OCT-SOX motifs on DNA. Taken together, we show that TOBF1 recruitment to DNA potentially mediated through lncRNAs like *Panct1* lead to rapid assembly of local nuclear regions (represented by puncta) where CTS events happen. This in turn orchestrates cellular events that maintain the fidelity of the mES cell pluripotent state.

**Extended Figure 4:**
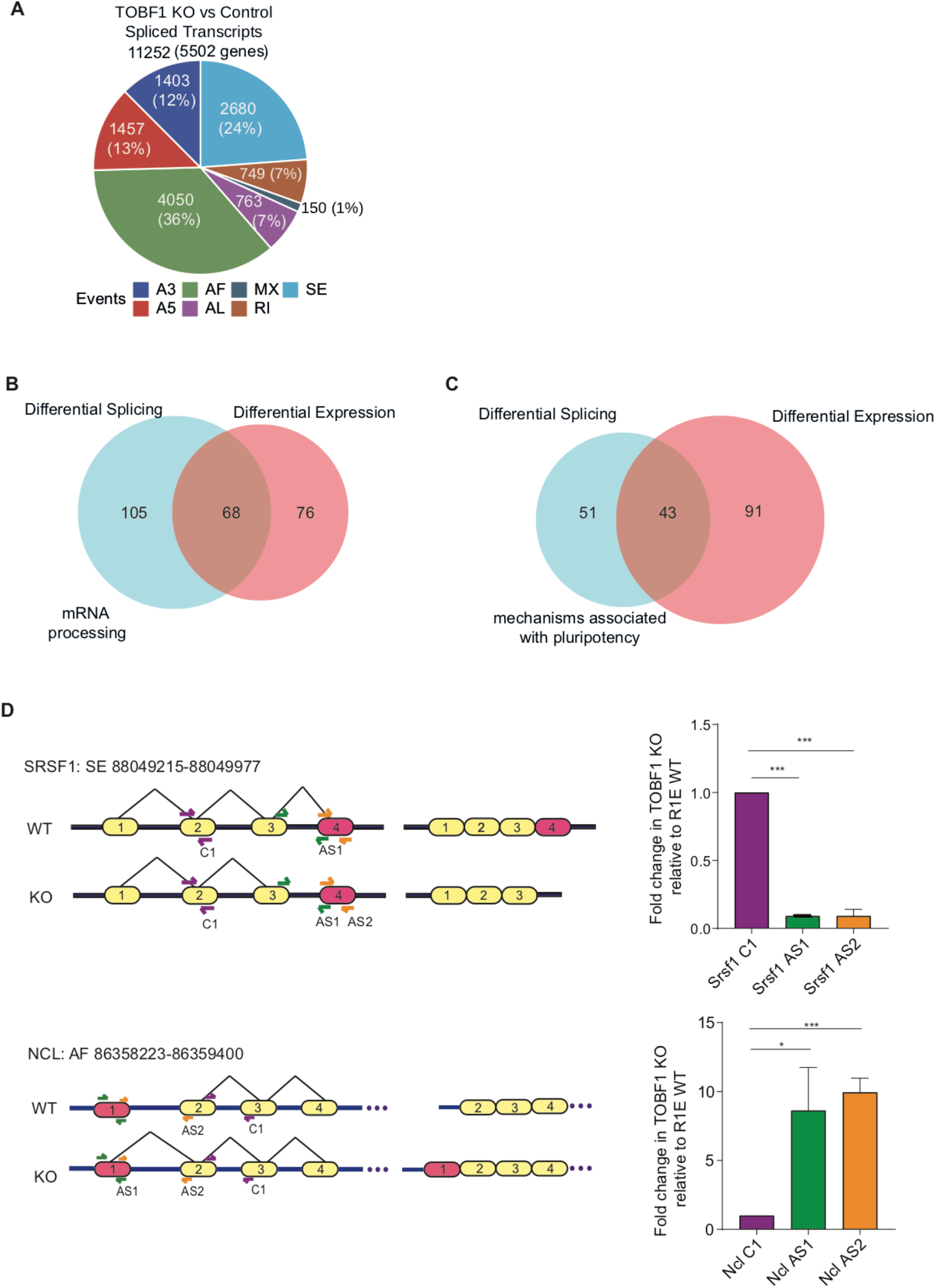
(A) Pie chart showing distribution of differential alternatively spliced events such as Alternative 3’-splice site (A3), Alternative first exon (AF), Mutually Exclusive (MX), Skipped Exon (SE), Alternative 5’-splice site (A5), Retained Intron (RI) (11252 transcripts out of 5502 genes) in TOBF1 KO and R1/E WT control cells. (B) Venn diagram showing overlapping transcripts count between differentially spliced transcripts and differentially expressed transcripts in mRNA processing pathways enriched in the gene ontology enrichment pathway analysis. (C) Venn diagram showing overlapping transcripts count between differentially spliced transcripts and differentially expressed transcripts in mechanisms associated with pluripotency pathways enriched in the gene ontology enrichment pathway analysis. (D) Schematic diagram showing the differences in the splicing pattern in Exon 4 of *Srsf1* and Exon 1 of *Ncl* gene transcripts isoforms in TOBF1 KO and R1E WT mES cells. qRT-PCR showing the relative mRNA fold changes using two different primer sets (AS1, AS2) to validate the splicing of Exon4 of *Srsf1* and Exon 1 of *Ncl* by taking constitutive exon as the control (C1) primer set. All values are normalized with β-Actin. Error bars indicate the SD of three independent replicates. *p<0.05, **p<0.001, ***p<0.0001 (Student’s t-test).

## Discussion

We propose a model (Figure 4C) whereby TOBF1 interacts with other mRNA processing and splicing factors to promote the alternative splicing of different transcripts essential for the regulation of ES cells. Its recruitment to chromatin at OCT-SOX motifs in early G1 phase suggests that TOBF1 may be a control switch for appropriate assembly of splicing factors and concomitant transcriptional regulation. Absence of TOBF1 resulted in the dissolution of such assemblies thereby affecting multiple downstream pathways. In our splicing and expression data we identified a myriad of ES cell specific genes, several of which function through independent pathways. Thus TOBF1’s role is upstream to canonical ES cell specific events although they cumulatively affect pluripotency.

It is interesting to speculate what the role of TOBF1 might be in other cell types. For example in mouse gastric epithelial cells, TOBF1 has been shown to suppress Wnt-related and Reg family genes ultimately leading to epithelial renewal. Indeed, in our TOBF1 RNA-seq data we did observe several genes involved in the Wnt pathway (105) to be differentially expressed (such as downregulated *c-Myc*, *Fosl1* and upregulated *Wnt7b, Fzd2* among others). However, our systematic study of chromatin association, cellular localization and CTS reveal a more detailed functionality of TOBF1 which expands beyond the scope of specific pathways and suggests a more elaborate central role of TOBF1 in regulating stem cell fate, at least in mES cells. Nevertheless, it will be important to investigate if TOBF1’s role in CTS is redundant in other cells, tissues and organs. A starting point to investigate this might be human ES cells where TOBF1 shows significant conservation.

In recent times, several groups have reported a strong association between lncRNAs and AS events with particular emphasis on their disease relevance. In several instances lncRNAs directly interact with splicing factors and cooperate or hijack AS events. Some examples include TPM1-AS lncRNA with RBM4 and DGCR5 lncRNA with SRSF1 both of which have roles in progression of human oesophageal cancer cells (106–108). Our studies highlight a similar cooperativity between the lncRNA *Panct1* and TOBF1 in the eventual formation of CTS hubs in mES cells. However, *Panct1* is poorly conserved across different species although TOBF1 shares significant homology between mouse, human, and other mammals. This suggests that additional lncRNAs might phenocopy *Panct1* in assembling such CTS factories. Also, as seen for other lncRNAs, local structural motifs such as g-quadruplexes or triplexes in RNAs might be the functional motifs linking such transcripts with CTS events (109–111). Systematic dissection of such motifs in *Panct1* and other TOBF1 interacting lncRNAs will shed light on how such cooperativity might occur.

A consistently intriguing observation in our study has been the bright G1 puncta where all these events occur. As these puncta bring together multiple proteins, DNA and RNA in close proximity, they suggest a dynamic assembly reminiscent of biomolecular liquid condensates. Our previous study showed TOBF1 puncta to exhibit dynamic coalescing properties akin to liquid-liquid phase separation. In a recent paper, the nuclear speckle marker SRRM2 has been shown to exhibit similar properties in HEK293T cells all across the cell cycle (46, 112). Since SRRM2 interacts physically with TOBF1, the TOBF1 puncta and associated CTS hubs in early G1 phase in mES cells possibly reflect phase separated condensates. Further studies through abrogation of defined TOBF1 motifs would be necessary to conclusively establish the identity of such condensates and the role of local protein-RNA contacts in maintaining them.

Although *Panct1* is very poorly conserved among vertebrates, TOBF1 shares a very high homology with multiple species underscoring its role in splicing and transcriptional regulation among others. Considering that it is largely uncharacterized to date, future studies might reveal non canonical roles that go over and beyond what we identified in this work.

## Methods

### mESC culturing and transfections

The mESCs were cultured in feeder-free complete media containing 1X DMEM (Gibco™, 11995-065), 20% Fetal Bovine Serum (Pansera Pan Biotech, P29-0705-ES), 1X Pen-strep (Gibco™, 15140122), 1% NEAA (Gibco™, 11140050), 0.1% Beta-Mercaptoethanol (Gibco™, 21985023) with additional supplement of 8ng/ml LIF (MPI-CBG, Dresden, Germany) per 500ml of media. The mESCs were cultured up to a maximum of 20 passages. Media were changed every day and split every alternate day by detaching with 0.25% Trypsin-EDTA (Gibco™, 25200056). Cells were incubated at 37 °C and 5% CO_2_.

For transfections, cells were seeded in 6 well or 12 well plates with 8×10^4^ or 4×10^4^ seeding density respectively. 16-20hrs post seeding, cells were transfected with 2ug per well (6 well plate) and 1ug per well (12 well plate) of plasmids concentration with the transfection reagent Lipofectamine 3000 (Invitrogen™, L3000001) and the protocol mentioned in the kit was followed. 48-72hrs post transfection, cells were harvested for downstream experiment.

For synchronizing ES cells in M phase, following trypsinization cells were washed in 1x PBS and released in fresh media containing 20 ng/ml Demecolcine (SigmaAldrich, D1925). After an incubation at 37 °C and 5% CO_2_ for 4h, cells were further washed twice with 1x PBS, released into Demecolcine-free media for 2h for entry into G1 and subsequently trypsinized and collected for further studies.

### Affinity pull down and mass spectrometry

Cell pellets were lysed in mild lysis buffer as previously described (Hubner et al., 2010 PMID: 20479470). The lysate was used as input material for affinity enrichment with magnetic beads precoupled to anti-GFP antibodies followed by trypsin digestion. The tryptic digest was desalted and analyzed on an EASY-nLC system (Thermo Scientific) connected to a Q-Exactive (Thermo Scientific) mass spectrometer (113). Maxquant software was used to analyze raw mass spectrometry data files (114). The data was filtered using the target-decoy strategy to allow a maximum of 1% false identification both on the peptide and protein levels.

### Generating TOBF1 KO cells

crRNAs targeting Exon 2, 3 and 4 (ChX: 59548256:159548275, 159551902:159551921, 159552068:159552087, 159553459:159553478) of Tobf1 gene were designed with CRISPOR tool (115). These crRNAs were cloned in SpCas9 with GFP reporter (Dual sgRNA Px458) vector which has scaffold for dual sgRNA cloning. This vector was designed for visualization and fluorescent based cell sorting. 4 crRNAs were designed and cloned using BbsI restriction enzyme based cloning strategy in 2 vectors using the protocol for sgRNA cloning (116) and confirmed with sanger sequencing. Oligo sequences of the crRNA forming DNA template and vector maps have been provided in the supplementary sheet. These two vectors were pooled and transfected with Lipofectamine3000 using the protocol provided in the kit. 48hrs post transfection, cells were harvested and single cells and bulk cells sorted based on GFP expression with BD FACSMelody™ cell sorter which uses the BD FACSChorus™ software. Cells are seeded in the same complete media and KO cells are confirmed with qPCR and Western Blot in the subsequent passages.

### Over-expression of *Tobf1*

Total mESCs RNA used to synthesize total cDNA with Superscript III First strand synthesis kit (Invitrogen™, 18080051). The full-length Tobf1 cDNA sequence (~2.3 kb) (UCSC ENSMUST00000112464.8 chrX:158331271-158375174) was then synthesized from the total cDNA by using Tobf1 specific cDNA AgeI containing forward and XhoI containing reverse primers which is listed in Supplementary table. The PCR was done by using Q5 High-fidelity DNA Polymerase (New England Biolabs, M0491) in a mix carrying 10 μl Q5 PCR buffer (5x), 2.5 μl 10 μM primers, 1ul 10mM dNTP mix (Invitrogen) and 5ul of total mESCs cDNA. The volume was adjusted to 50 μl using PCR grade water. The reaction was set up in a DNA Thermocycler (Bio-Rad) and the conditions for PCR were: Initial denaturation at 98 °C for 2 min followed by 40 cycles of denaturation, annealing and extension (98 °C for 10 secs, 65 °C for 45 secs, 72 °C for 3 mins) and a final extension at 72 °C for 10 mins.

The PCR amplicons were purified using a PCR purification kit (Qiagen). This PCR amplicon was fused with Kozak 3XHA (~100 bps) amplified product by using overlapping primers having NheI forward and XhoI containing primers and the PCR was conducted using the same protocol mentioned above. This 3XHA Tobf1 CDS product (~2.4 kb) was digested with NheI and XhoI restriction enzymes and ligated into the vector pCAGGS-RFP in which the RFP sequence was replaced with the PCR product. Successful cloning was confirmed by Sanger Sequencing.

### Bulk RNA sequencing

#### RNA extraction methods

For extracting cells in the G1 phase, ES FUCCI vector was transfected in TOBF1 KO, OE and Control cells and as G1 phase is indicated by the expression of reporter gene mCherry, we FACS sorted cells expressing either mCherry (G1) or non-mCherry (non G1) cells and examined the steady state levels of Tobf1 in these two sorted populations. For RNA isolation, TOBF1 over-expressed and knock-out samples along with controls are sequenced in three independent biological replicates on Illumina Hi-Seq platform.

### RNA sequencing analysis

Quality control for all samples was assessed using FastQC. Low-quality reads and adaptor contamination were removed through Trimmomatic. The RSEM reference index was generated using mouse reference genome (GRCm39/mm39) and transcriptome (gencode.vM27) with the rsem-prepare-reference command coupled with STAR aligner. Subsequently, Gene expressions were estimated by alignment of sequencing reads to the reference index with STAR and rsem-calculate-expression command. Further, gene count matrix was generated with test and control samples for both the knockout and overexpression of Tobf1. Gene expression counts were normalized using TMM followed by differential expression using generalized linear model quasi-likelihood F-tests (glmQLFTest) from the edgeR Bioconductor package. Top differentially expressed genes were visualized with the help of the EnhancedVolcano Bioconductor package. The Complex Heatmap package from Bioconductor was used to generate heatmap.

For estimating the splicing events, a pipeline, as described in the SUPPA2 package, was executed. As recommended in the SUPPA2 tutorial, reference indexes were built from mouse transcriptome (gencode.vM27) followed by the pseudo alignment of the reads to generate transcript counts using salmon aligner. Further, reference for various splicing events is built from gencode.vM27 using ‘generateEvents’ sub-command of SUPPA2. Transcript quantification files from the salmon aligner were used to generate a matrix followed by the estimation of local alternative splicing events with ‘psiPerEvent’ subcommand of SUPPA2. Differential splicing events were calculated by diffSplice SUPPA2 sub-command with the empirical method. Any splicing events with a difference of greater than |0.10| PSI were considered statistically significant. Splicing events between different conditions are visualized using sashimi plots with ggsashimi tool. Gene expression, splicing and ChIP-Seq data is integrated and visualized through the Circilize package.

For pathway and gene enrichment analysis, clusterProfiler has been fed with differential expressed transcripts to predict the gene ontology and wiki pathways. This process is also performed for genes with differential splicing events as well as for proteomics data from mass-spectroscopy.

### qRT-PCR

Total RNA was isolated from mES cells 48 hrs post transfection. Qiagen RNeasy kit (74106) was used to isolate total RNA in the step-wise protocol provided with the kit. The RNA isolated was treated with DNaseI (Turbo DNase kit, Invitrogen™, AM2238). cDNA synthesis was done from DNaseI treated total RNA with Qiagen cDNA synthesis kit (205313) with the incubation conditions suggested with the kit. cDNA synthesis was followed by real time qPCR for the test and control samples in triplicates. The transcripts were quantified by using SYBR Green Master Mix: SYBR green based TB Green Premium Ex Taq II (Takara, RR82WR) in the PCR instrument Light Cycler 480 (Roche) or BioRad. All the Ct values obtained for different transcripts were normalized with the Ct value of Beta-Actin. The Fold change analysis of the transcripts for comparative analysis was done using the 2^-ΔΔCt^ method (117)

### Western Blot

The ES cells were lysed using RIPA lysis buffer (ThermoFisher, Pierce™) to prepare the cell protein lysate. 100 μl of the lysis buffer was added to each 6-well plate along with 1×Protease Inhibitor Cocktail (PIC, Roche cOmplete). The cells were incubated in a rocker at 4°C for 1hr. Protein lysate from each sample was collected and the concentration of the protein was estimated using Pierce™ BCA Protein Assay Kit (ThermoFisher). For each sample, 30 μg of protein was loaded into the wells of 10% SDS gel and PAGE was performed using SDS running buffer (2.5mM Tris base, 19mM Glycine, 0.1% SDS in autoclaved milliQ). The proteins were then transferred from the gel to the PVDF membrane (GE Healthcare Life-Science) in Bio-Rad vertical gel Transfer Apparatus using Transfer buffer (2.5mM Tris base, 19mM Glycine, 20% v/v Methanol in autoclaved milliQ) at 4°C for 1.5 hrs at 95 V. After the transfer was complete, the membrane was cut according to the required protein size and kept for blocking with 5% BSA in 1×TBST (20mM Tris base, 150 mM NaCl and 0.2% Tween-20) on a rocker at room temperature for 2 hr. After blocking, the blots were incubated with primary antibody (listed in the supplementary table) at 1:1000 dilution in the same blocking buffer overnight in a rocker at 4°C. Beta-Tubulin antibody (1:5000 dilution) is taken as the loading control. After primary antibody incubation, the blots were washed three times for 10 min each with 0.2% TBST. Post washing, the same blots were incubated with a secondary antibody having HRP conjugate in a rocker for 2 hr at room temperature. Post secondary antibody incubation, the blots were washed three times for 15 mins each. For signal development, EMD Millipore™ Immobilon Western Chemiluminescent HRP Substrate (ECL) was used to develop the blots in the Syngene Gel doc instrument. ImageJ was used for the densitometry analysis and quantification of the signals.

### Immunofluorescence (IF)

For slide preparation, R1E mES cells or TOBF1 GFP cells were seeded in 0.1% Gelatin pre-coated 22×22mm coverslips (Corning, CLS285022) placed in a 6 well plate. ~8×10^4^ cells seeded and 24hrs post seeding and adherence, cells were harvested by washing with 1×PBS (Gibco™, 10010023) twice in a well plate. All the procedure since harvesting was carried out in the well plates that contained the cell adhered coverslips. Cells were then fixed with chilled 4% Paraformaldehyde (PFA) + EDTA (Ph 7.4) by incubating at Room Temperature (RT) for 10 mins in a mild shaker. Post fixation, the coverslips were washed twice with a washing buffer (10μM MgCl_2_, 5μM EGTA in 1×PBS). Cells were then incubated with a permeabilization buffer (0.2% Triton X-100, 10μM MgCl_2_, 5μM EGTA in 1×PBS) at RT for 10 mins followed by washing with a washing buffer twice. After permeabilization, cells were incubated with blocking buffer (2% BSA, 10μM MgCl_2_, 5μM EGTA in 1×PBS) for 15 mins at 37°C. After blocking, the cells were incubated with primary antibodies (listed in supplementary sheet) in the ratio of 1:250 dilution in the same blocking buffer. Incubation was done for 2hrs at 37°C or overnight at 4°C. For Co-IF studies, two antibodies raised in different species were mixed in 1:250 dilution ratio in blocking buffer and incubated in the same manner post fixation and permeabilization of the cells in coverslips. Once the primary incubation time was completed, the cells were washed with 1×PBS thrice for 5mins each. Now the cells were incubated with Alexa fluor labeled fluorescent secondary antibodies (ThermoFisher Scientific) to be conjugated specifically with the primary antibody, considering the species in which it is raised. Fluorescently tagged secondary antibodies were used in the dilution ratio of 1:1000 in the same blocking buffer and incubated for 30 mins at RT. Post incubation, the cells are washed with 1×PBS thrice for 5mins each. After the wash, one drop of ProLong™ Diamond Antifade mountant with DAPI (Invitrogen™, P369366) was added in each coverslip and these coverslips were then mounted on one side frosted glass slides (Corning, CLS294875X25) and proceeded for imaging. DeltaVision™ Ultra-light microscope (GE Healthcare) was used for imaging the slides at 60X objective with the help of DV SoftWorx software. Co-localization quantification studies of all images were done in ImageJ with Coloc2 and JACoP plugins by determining the Pearson’s and Mander’s coefficients.

### Immunoprecipitation (IP)

1×10^6^ cells were seeded in a 100mm dish (Corning) and 16hrs post seeding appropriate transfection was done according to desired experiment. 48hrs post transfection, cells were washed and trypsinized and cross-linked with 1% Formaldehyde by incubating at RT for 15mins. Formaldehyde was quenched by adding 125mM Glycine to the formaldehyde and incubated at RT for 10mins. Pellet down the cells by centrifuging at 1200 rpm for 5mins. The cell pellet was washed twice and lyse the cells with RIPA cell lysis and extraction buffer (Thermo Scientific™, 89900) and 1x Protease Inhibitor cocktail (PIC) (Roche, SKU11697498001). The total protein concentration in the cell lysate was quantified by using BCA protein Assay kit (Thermo Scientific™, 23225).

Pre-clearing of lysate: Equal concentration of lysate taken for test and control samples and these lysates were pre-cleared with 5ug of IgG isotype control (ab171870) and 15ul of 30mg/ml Dynabeads protein A (Invitrogen™, 10001D) at 4°C for 2hrs on a rotator.

#### Beads blocking

In separate vials, 35ul of Dynabeads Protein A is taken for each sample for immunoprecipitation and these beads are once washed with 1×PBS and then allowed to incubate with 2% BSA and 10ng/ml Yeast tRNA (Thermo Scientific™, AM7119) for blocking the beads to avoid non-specific binding of proteins and RNAs. This incubation is done at 4°C for 2hrs on a rotator.

#### Antibody binding with blocked beads

Blocked Beads are incubated with 5-10ug of ChIP grade anti-HA antibody (ab9110) in 1×PBS and allowed to incubate at 4°C for 4hrs on a rotator.

#### Lysate and beads binding

Beads from Pre-cleared lysate is removed by placing it in a magnetic stand and this pre-cleared lysate is then allowed to incubate with the antibody bound beads 4°C for 12-16hrs (O/N) on a rotator. A small volume of each sample is kept aside as 1-5% input which is not subjected to antibody binding and pull-down.

After incubation completion, the beads samples were washed with RIPA buffer thrice and then the beads were incubated with Elution buffer (1%SDS, 10mM NaHCO_3_) at 30°C for 30mins. The eluted samples were checked with Western blot along with 1% input sample.

### Cell cycle assay

The cells are harvested by trypsinizing them and washing them with cold 1×PBS. This was spinned at 1500xg for 5 mins. The pellet was fixed with chilled 70% Ethanol in a dropwise manner, while gently vortexing the pellet. This was again spinned at 1500 xg for 5 mins. The pellet was resuspended with a buffer (0.1% Triton-X+RNaseA in 1×PBS) and incubated at 37°C for 3 hrs. Post incubation, Propidium Iodide (PI) (5ug/ml) was added to all the samples required for acquisition. For acquisition of the samples, BD LSR II FACS system was used and analysis of different cell cycle was done using FlowJo v10.6 software.

### Statistical analysis

The statistical analysis for experiments was performed using GraphPad Prism 8.4 to evaluate significance among experimental replicates. All data were presented as mean ± S.D. of three independent biological replicates except otherwise mentioned. A two-tailed unpaired Student’s *t*-test was used to analyze the q-RT-PCR and WB experimental data. The experimental results leading to a *P*-value <0.05 were considered statistically significant. One asterisk (*), two asterisks (**), three asterisks (***) denote *P* < 0.05, *P* < 0.005 and *P* < 0.0005, respectively.

## Data availability

The TOBF1 KO RNA sequencing files are submitted in the Sequence Read Archive (SRA) database with the Project ID: **PRJNA900687.** Reviewer link is here: https://dataview.ncbi.nlm.nih.gov/object/PRJNA900687?reviewer=u7k2as985aipu5h81l4laigb4m

## Supplementary files information

**Supplementary file 1:** Total differentially enriched interacting proteins observed in TOBF1-GFP pull down mass spectrometry. Sheet 1: Based on the p-values and peptide identifications in all the control measurements, yellow marked proteins are a very convincing interactor as they are never seen in any control. Green marked proteins are very likely interactors which were seen only once in control experiments in much lower intensities. Sheet 2: List of true potential interactors. Sheet 3: List of proteins from the interactome profile that are involved in mRNA processing and splicing

**Supplementary file 2:** Differential gene expression in total RNA sequencing of TOBF1 KO and R1/E WT control. Sheet 1: Gene expression count. Sheet 2: Differential expressed genes with fdr <0.05.

**Supplementary file 3:** Pathway analysis of differentially expressed transcripts: Sheet 1: Biological processes. Sheet 2: Molecular functions. Sheet 3: Kegg pathways. Sheet 4: Reactome. Sheet 5: Phenome data. Sheet 6: Wikipathways

**Supplementary file 4:** Differential alternatively spliced events of global transcript isoforms in total RNA sequencing of TOBF1 KO and R1/E WT control. Differential spliced events of various transcripts with Dpsi |0.10|

**Supplementary file 5:** Pathway analysis of differentially spliced transcript isoforms: Sheet 1: Biological processes. Sheet 2: Molecular functions. Sheet 3: Kegg pathways. Sheet 4: Reactome. Sheet 5: Phenome data. Sheet 6: Wikipathways

## Acknowledgments

The authors thank Dr Amrita Singh and Ms. Rhythm Phutela from CSIR-IGIB for designing and creating dual sgRNA SpCas9 plasmid and dFnCas9-KRAB plasmid, respectively which is used for TOBF1 Knockout and CRISPRi experiments. The authors are grateful to all members of the Chakraborty and Maiti labs for helpful discussions and suggestions and Mr. Manish Kumar, Imaging Facility CSIR IGIB for help with microscopy experiments. This study was funded by an EMBO Young Investigator Award (GAP0252) and Department of Biotechnology grant (GAP0188) to D.C. The MS experiments were funded by EU FP7 grant SyBoSS (242129) to F.B.

## Contributions

M.A., S.M. and D.C. conceived, designed and interpreted the experiments. A.H.A analyzed and provided bioinformatics support for data interpretation. L.D. and F.B. analyzed and interpreted data from BAC tagged cell lines. V.I. and C.C. performed and analyzed data from affinity purification studies. D.P. performed experiments with FUCCI tagged ES lines. M.A. and D.C. wrote the manuscript with inputs from all other authors.

